# Cytoplasmic expression of the cell cycle regulator cyclin D1 in radial glial progenitor cells modulates brain cortex development

**DOI:** 10.1101/2024.12.23.630056

**Authors:** Neus Pedraza, Daniel Rocandio, Maria Ventura Monserrat, Bahira Zammou, Ariadna Ortiz-Brugués, Pau Marfull-Oromí, Disha Chauhan, Mario Encinas, Xavier Dolcet, Carme Espinet, Francisco Ferrezuelo, Eloi Garí, Joaquim Egea

## Abstract

During nervous system development, the interplay between cell cycle regulation and neurogenesis is fundamental to achieve the correct timing for neuronal differentiation. However, the molecular players regulating this transition are poorly understood. Among these, the cell-cycle regulatory cyclins and their cyclin-dependent kinases (Cdks) play a pivotal role. In the present work we uncover an unknown function of cyclin D1 (Ccnd1) during cortex development which is independent of cell cycle regulation and that relies on its cytoplasmic localization and membrane association. We show that Ccnd1 is localized in the cytoplasm of the radial glial process (RGP) of neuron progenitors in different regions of the developing brain, including the cortex. Cytoplasmic Ccnd1 is enriched at the distal tip of the RGP, adjacent to the meningeal basement membrane, and overlaps with β1-integrin at the plasma membrane. *CCND1* knock-out animals show an abnormal cortical layering in which the distribution of Tbr2+ and Ctip2+ cells are affected without displaying proliferation defects. This is consistent with a cytoplasmic function of Ccnd1 as overexpression by in utero electroporation of a dominant negative Ccnd1, unable to activate Cdks, and targeted to the cytoplasmic membranes, reproduces some of these Tbr2 and Ctip2 defects. Finally, we provide evidence that cytoplasmic Ccnd1 affects neuron morphology and that it is required for the proper detachment of the RGP from the meningeal basement membrane by a mechanism involving the phosphorylation of the integrin effector protein paxillin. Hence, we propose that Ccnd1 has an important cytoplasmic function for cortical development independently of cell cycle regulation.

**Significant Statement:** A key developmental step during nervous system formation is the transition from proliferating progenitors to postmitotic neurons. However, the molecular mechanisms regulating this process are not fully understood. Cyclin D1 (Ccnd1) is a canonical regulator of cell cycle in the cell nucleus. Surprisingly, we show that Ccnd1 is also located in the radial glial process of neuron progenitors and associated to the plasma membrane in different regions of the developing mouse brain. We uncover a novel function for this cytoplasmic Ccnd1 and show that it is required for proper cortical layering, independent of cell cycle regulation. Mechanistically, we provide evidence that this function is mediated by the integrin effector paxillin. We propose therefore that cytoplasmic Ccnd1 is important for cortex development independent of cell cycle regulation.

## Introduction

Projection neurons within the cerebral cortex are organized in a six-layered structure, each one with specific molecular identities and functions. During development, these neurons differentiate from proliferative progenitor cells in the ventricular zone (VZ, apical progenitors) or in the subventricular zone (SVZ, basal progenitors). Cell cycle regulation plays an important role during neuron differentiation. Symmetric divisions give rise to two daughter cells that maintain the progenitor status whereas in asymmetric, neurogenic divisions, one of the daughter cells exits the cell cycle and differentiates into a postmitotic neuron which finally migrates towards the cortical plate (CP) and populates a specific cortical layer (Delaunay et al., 2017). Proliferative versus neurogenic divisions are determined by the duration of the G1 phase of the cell cycle so that the lengthening of the G1 phase stimulates neurogenesis by promoting neurogenic divisions (Takahashi et al., 1995; Lukaszewicz et al., 2005; Pilaz et al., 2009). However, the exact mechanisms that control the balance between cell renewal and lineage commitment are still poorly understood.

D-type cyclins (Ccnd1-3) control cell cycle length by acting as regulatory subunits of cyclin-dependent kinases (Cdks) 4 and 6. Ccnd/Cdk complexes inhibit the pRB repressor by phosphorylation, releasing the E2F transcription factor and promoting the expression of genes necessary for cell cycle entry (Sherr et al., 1993). Overexpression of Ccnd1/Cdk4 promotes G1 shortening, expansion of neural progenitors and a delay of neurogenesis (Lange et al., 2009; Artegiani et al., 2011), and alterations in Ccnd1 levels in neural progenitors have been related to cortical defects in Down syndrome (Najas et al., 2015). These results are in contrast with in vivo data in mice. *CCND1* knock-out animals do not display proliferation defects in the neocortex (Glickstein et al., 2009), and just mild defects in the cerebellum that lead to a reduction of cerebellar size (Pogoriler et al., 2006). Nevertheless, deficient *CCND1* mice show symptoms of neurological impairment as indicated by an abnormal hindlimb clasping reflex, although the underlying causes have not yet been established (Sicinski et al., 1995).

Non-canonical functions of Ccns/Cdks have been reported (Hydbring et al., 2016). Ccnd1/Cdks can directly regulate the activity of transcription factors such as ATOH1 or Hes6 and control neuronal commitment in a cell-cycle independent manner (Miyashita et al., 2021; Lukaszewicz and Anderson, 2011). Interestingly, Ccnd1 has also been reported to be associated to the plasma membrane and regulate cytoplasmic targets such as β1-integrin and its downstream effector paxillin and affects cell migration and cell-matrix adhesion (Fusté et al., 2016a; Fernández-Hernández et al., 2013). Cytoplasmic-associated Ccnd1 activity is also relevant for neural communication as it regulates gamma-aminobutyric acid (GABA) signaling in rat hippocampal neurons (Pedraza et al., 2023).

In the present work, we demonstrate that Ccnd1 is expressed in the cytoplasm of radial glial cells in the developing cortex overlapping with β1-integrin at the plasma membrane of the tip of the radial glial process (RGP). *CCND1* knock-out animals display an abnormal cortical layering of Tbr2+ and Ctip2+ neurons but this phenotype is not accompanied by proliferation effects. Instead, some aspects of these cortical layering defects are reproduced when a cytoplasmic dominant negative Ccnd1 is overexpressed by in utero electroporation, suggesting a function for cytoplasmic Ccnd1 in cortical layering. Mechanistically, we suggest that this cytoplasmic Ccnd1 affects neuron morphology both in vivo and in vitro, and that it is important for the proper detachment of the RGP from the meningeal basement membrane through the phosphorylation of the integrin signaling complex effector, paxillin. Therefore, we propose that Ccnd1 has an important cytoplasmic function in cortex development that is separate from its role in regulating the cell cycle.

## Materials and Methods

### MICE

Animal care followed the Guidelines of the University of Lleida for Animal Experimentation in accordance with Catalan, Spanish, and European Union regulations (Decret 214/1997, Real Decreto 53/2013, and Directive 63/2010). Animals were housed in the animal care facility of the University of Lleida with 12:12h light/dark cycle and food/water available ad libitum. *CCND1* knock-out mice (Sicinski et al., 1995) were obtained from Charles-River and backcrossed to C57/bl6 background. For embryo dissection, the day on which the vaginal plug was found was considered as embryonic (E) day 0.5 and the date of birth was considered as postnatal (P) day 0. PCR genotyping was done by tail biopsy using the following primers: common primer CTGTCGGCGCAGTAGCAGAGAGCTACAGAC, *CCND1* wild-type allele specific primer CGCACAGGTCTCCTCCGTCTTGAGCATGGC and a *CCND1* knock-out specific primer CTAGTGAGACGTGCTACTTCCATTTGTCACG. The expected band sizes are 249 bp for the WT allele and 394 bp for the *CCND1* knock-out allele.

### EXPRESSION VECTORS

One copy of His tag or three copies of the HA epitope were added at 5’ end of the human *CCND1* or mouse *CDK4* open reading frames (ORF). *CCND1^CAAX^* or *CDK4^CAAX^* constructs are fusions of the 3′ end of the K-Ras ORF containing the CAAX motif (GGC TGT GTG AAA ATT AAA AAA TGC ATT ATA ATG TAA) at the 3′ end of the *CCND1* or *CDK4* ORFs (Fusté at al, 2016b). Standard PCR-mediated site-directed mutagenesis was used to obtain the K112E mutant of *CCND1* (Fernández-Hernández et al., 2013). Mouse wild-type and non-phosphorylatable paxillin were previously described (Fusté et al., 2016a). Constructs were subcloned into the expression vectors pcDNA3 or pCAGIC (Addgene, 11159). pcDNA3 was used for neuron transfection in vitro and was co-transfected with pEGFP-N1 plasmid (Clontech) to assess neuron morphology. pCAGIG contains an internal ribosome entry site (IRES) and the EGFP gene for tracking electroporated cells in the in utero electroporation experiments (IUE).

### INTRAVENTRICULAR INJECTION AND IN UTERO ELECTROPORATION

E13.5 pregnant CD1 wild-type mice were deeply anesthetized with isoflurane (IsoFlo, Zoetis) during the entire operation procedure. To relax uterus muscles, β2 agonist Ritodrine (Sigma R0758) was administered intraperitoneally, and buprenorphine (Buprex, 100 mg/ml) subcutaneously as an analgesic. A 2 cm laparotomy section was made in the abdomen, and the uterine horns were carefully exposed and lubricated with NaCl 0.9% at 37° C. Two to 4 microliters of purified plasmid DNA dissolved in PBS (1 μg/μl) containing 0.025% of Fast Green (Sigma-Aldrich) was injected in the lateral ventricles of each embryo using a glass capillary (World Precision Instruments) sharpened previously by Puller P-97 (Sutter Instrument). Platinum electrodes (CUY701P20L, Nepagene) were placed across the head with the positive pole adjacent to the neocortex, to enhance the permeability of the cell membrane and allow the entrance of DNA. Five 30 mV electric pulses of 50 ms with intervals of 950 ms were charged by an electroporator (ECM830, BTX). Uterine horns were placed back into the abdominal cavity and abdomen wall and skin were surgically sutured. During the whole operation embryos were manipulated with ring forceps (Fine Science Tools). Embryonic brains were dissected, as indicated, at different time-points after electroporation and fixed directly in ice cold 4% paraformaldehyde (PFA) solution in PBS overnight. Postnatal brains were obtained after intracardial perfusion with ice-cold PBS and ice-cold 4% PFA-PBS and pos-fixed overnight in ice-cold 4% PFA-PBS.

### EdU LABELING

E14.5 pregnant mice were injected intraperitoneally with 100 μg/g body weight of EdU (ThermoFisher, C10337) using a stock solution of 10 mg/ml in PBS. Pregnant mice were sacrificed 1h later and the embryonic brains were processed for cryosectioning (see below) and immunostaining with anti-EdU antibody. Sections were washed with PBS and permeabilized and blocked with 5% goat serum in 0.1% Triton X-100 in PBS for 1 h at room temperature. Subsequently, the slides were incubated during 30 minutes with Click-iT reaction cocktail (ThermoFisher, C10337) protected from light. After washing, DAPI was added to sections during 2 h at room temperature for nuclei staining.

### CORTICAL NEURON CULTURE AND NEURITE ANALYSIS

Primary culture of cortical neurons from E15.5 *CCND1* knock-out, control littermates or wild-type embryos were prepared as previously described (Pedraza et al., 2023). To identify the different genotypes, a PCR reaction was set-up during the dissection procedure using a tail biopsy as indicated above. Primary cortical neurons were electroporated using Ingenio electroporation solution (Mirus) following the manufacturer’s recommendations and the proportion 2 μg DNA:2*10^6^ cells. The media was changed 2 hours after transfection and cells were fixed at 1-4 days in vitro (DIV). For Ccnd1 downregulation, scrambled shRNA (SHC002, Sigma) or Ccnd1 shRNA (TRCN0000026883, Sigma) were co-transfected with EGFP-N1 in a proportion 2:1, by in-tube transfection as described previously (Halterman et al., 2009) with Lipofectamine 2000 (ThermoFisher). For neurite analysis, cortical neurons were fixed using 4% PFA and 4% sucrose in PBS for 15 min at room temperature and then washed with PBS.

Neurons were permeabilized for 5 min with 0.1% Triton X-100 in PBS and blocked with 3% BSA in PBS. Primary antibodies α-GFP (ThermoFisher, A11120), α-cleaved caspase-3 (Cell Signaling, 9661) or α-HA (Sigma, 3F10) were diluted 1:200 in 0.3% BSA-PBS. Proteins were detected by incubation with fluorescent-labelled secondary antibodies (ThermoFisher). Hoechst (Sigma) was used to label the nuclei. Immunofluorescence images were obtained with an Olympus IX71 microscope and quantification of neurite length was performed using NeuronJ of ImageJ. All the analysis were done blind to the experimental condition.

### LENTIVIRAL PRODUCTION AND MEF INFECTION

For lentivirus production, HEK293T cells were transfected with lentiviral expression vectors, envelope plasmid pVSV.G, and packaging plasmid pHR’82ΔR at a ratio of 2:1:1. For RNA interference, the ccnd1 MISSION shRNA TRCN0000026883 and the control SHC002 cloned in a pLKO.1-puro were obtained from Sigma-Aldrich. MEFs were infected and selected with Puromycin.

### TISSUE IMMUNOFLUORESCENCE AND ANTIBODIES

Embryonic and postnatal brain tissue were processed, cryosectioned and immunostained as previously described (Fleitas et al., 2021). Antibodies used were: goat α-GFP (Abcam, ab6673; 1:300), rabbit α-Ccnd1 (Dako, M3642; 1:300), rabbit α-Tbr2 (Abcam, ab23345; 1:300), rat α-phospho-histoneH3 (PH3) (Sigma-Aldrich, H9908; 1:100), rabbit α-Sox2 (Abcam, ab97959; 1:300), rabbit α-Tbr1 (Abcam, ab31940; 1:300), rat α-Ctip2 (Abcam, ab18465; 1:300), rabbit α-Cux1 (Proteintech, 11733-1-AP; 1:200), mouse α-Pax6 (hybridoma bank, 1:300), mouse α-nestin (Abcam, ab6142; 1:100), mouse α-βIII-tubulin (Sigma-Aldrich, T8578), α-β1-integrin (Chemicon Millipore, MAB1997). Fluorescent-labelled secondary antibodies were from Jackson ImmunoResearch and were used at 1:300.

### IMMUNOBLOTTING

*CCND1* knock-out protein was extracted from embryo hands in 0.83M Urea, 0.105M Tris-HCl pH6.8 and 1.7% SDS, with glass beads in a bead beater homogenizer. For immunoblot, protein samples in Laemli buffer with 1% β-mercaptoethanol were resolved by SDS-PAGE, transferred to PVDF membranes (Millipore), and incubated with primary antibodies: α-Ccnd1 (monoclonal DCS-6, BD biosciences, 556470; 1:500), α-Ccnd1 (polyclonal antibody ABE52, Millipore, 1:1000), α-actin (monoclonal C4, Millipore), α-Pax (monoclonal, BD transduction, 610051, 1:1000), α-pS83 Pax (polyclonal, ECM Biosciences, PP1341, 1:500), α-pS178 Pax (polyclonal, Calbiochem, ST1069, 1:500). Appropriate peroxidase-linked secondary antibodies (GE Healthcare UK Ltd) were detected using the chemiluminescent HRP substrate Immobilon Western (Millipore). Chemiluminescence was recorded with a ChemiDoc-MP imaging system (BioRad).

### RNA ISOLATION AND REAL-TIME QUANTITATIVE PCR

RNA from cultured cortical neurons collected at different DIV was isolated with RNeasy mini kit (Qiagen), treated with DNAseI (Sigma) and retrotranscribed with SuperScript Reverse Transcriptase (Thermofisher). The obtained cDNA samples were analyzed by Real-time quantitative PCR in a reaction containing 25ng of cDNA, 1xSYBR Green master mix (Applied Biosystems™ SYBR™, Thermofisher) together with forward and reverse primers (150 nM) for Ccnd1 (forward 5’-GCGTACCCTGACACCAATCTC-3’ and reverse 5’-CTCCTCTTCGCACTTCTGCTC-3’) or GAPDH as a housekeeping control (forward 5’-AGGTCGGTGTGAACGGATTTG-3’ and reverse 5’-TGTAGACCATGTAGTTGAGGTCA-3’). qPCR was performed in the C1000 Thermal Cycler CFX96 Real-Time System (Bio-Rad Laboratories) with the following conditions: initial denaturation at 95°C for 2 min; 40 cycles at 95°C for 15 s, 65°C for 1 min.

### STATISTICAL ANALYSIS

To evaluate two experimental groups, two-tailed unpaired Student’s T test (*t*-Student) was performed. For multiple comparison, one-way ANOVA with a Tukey post-hoc analysis or two-way ANOVA with Sidak’s post-hoc analysis was performed. Microsoft Excel and GraphPad Prism v5.0 were used for statistical analysis and graphical representation. The *p* value in each experiment is indicated, and significance was considered when p<0.05 (*), p<0.01 (**) or p<0.001 (***). Error bars were calculated using the standard error of the mean (SEM).

## Results

### Expression pattern of Ccnd1 in the developing brain

To understand the function of Ccnd1 during nervous system development we studied its expression in the brain by immunofluorescence with specific antibodies. As expected, we observed a typical nuclear localization of Ccnd1 in proliferative regions (VZ and SVZ) alongside the ventricle of embryonic brains, including the cortex, basal ganglia and thalamus (Fig. 1A). Interestingly, in the cortex we noticed that Ccnd1 displayed a graded expression, from ventro/lateral-high to dorso/medial-low (Fig. 1A,D), that contrasted with the homogenous expression of other proliferation markers such as Sox2 or anti-phospho histoneH3 (PH3) (Fig. 1B,C,E,F)(Borello et al., 2018). Surprisingly, Ccnd1 staining revealed fiber-like structures in the cortex, thalamus and basal ganglia indicating a cytoplasmic localization of the protein (Fig. 1A,G,H,I). Nuclear and cytoplasmic Ccnd1 stainings were specific since both disappeared when *CCND1* knock-out tissue was used (Fig. 1J-L). Fiber-like Ccnd1 localization co-stained with Nestin but not with βIII-tubulin, indicating the presence of cytoplasmic Ccnd1 specifically in progenitor cells (Fig. 1M-R and data not shown). A closer look at this staining showed that Ccnd1 localization at the RGP was enriched at the distal, basal end, forming a button-like structure adjacent to the meningeal basement membrane which overlaps partially with the adhesion protein β1-integrin at the plasma membrane (Fig. 1M,P,S-V). In summary, we have described three specific and distinct Ccnd1 localizations in the mouse developing brain: i) nuclear, in the VZ/SVZ, with a graded pattern from ventro/lateral-high to dorso/medial-low; ii) cytoplasmic at RGPs and iii) at the tip of basal end of the RGP forming a cytoplasmic button-like structure, overlapping with β1-integrin at the plasma membrane, adjacent to the meningeal basement membrane. Association of Ccnd1 with the plasma membrane was previously reported in non-neuronal cells such as keratinocytes, fibroblasts and cancer cells (Zhong et al., 2010; Fernández-Hernández et al., 2013; Fusté et al., 2016a; Fusté et al., 2016b).

**Figure 1.**
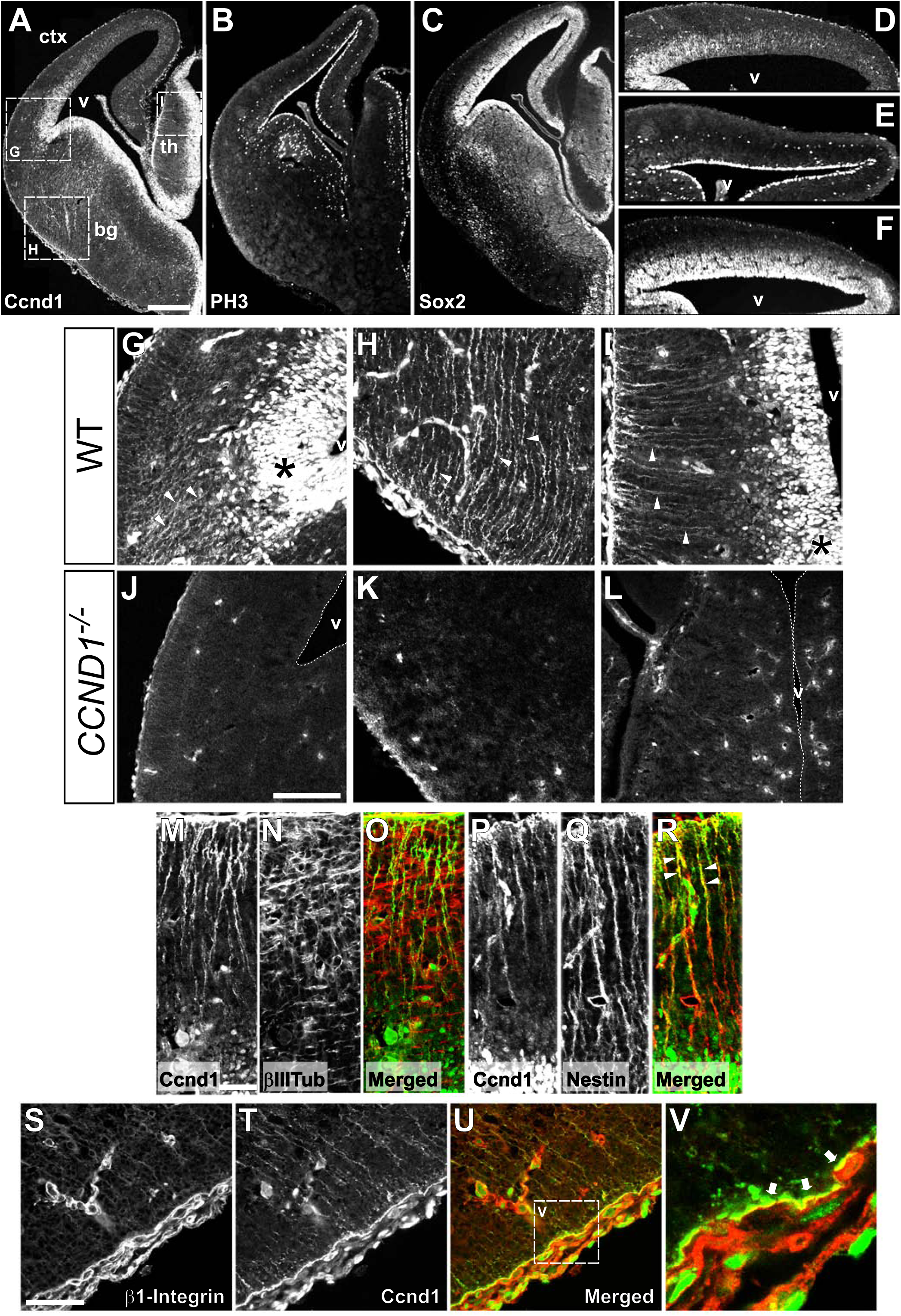
Expression analysis of Ccnd1 during brain development reveals a cytoplasmic localization of the protein in the RGPs. A-C) Images of cryosections of wild-type embryonic brains at E14.5 (only one half is shown) stained by immunofluorescence with antibodies against Ccnd1 (A), PH3 (B) or Sox2 (C). Dashed squares in A indicate the position of panels G-I. D-F) Magnified images of the cortex shown in A-C, stained by immunofluorescence with antibodies against Ccnd1, PH3 or Sox2. G-I) Magnified images of the regions shown in A of the pallial-subpallial boundary (G), basal ganglia (H) or thalamus (I) stained by immunofluorescence with antibodies against Ccnd1. Asterisks indicate the nuclear staining in the progenitor cell niche. Arrowheads point to the fiber-like staining. J-L) Images of anti-Ccnd1 immunofluorescence of the pallial-subpallial boundary (J), basal ganglia (K) or thalamus (L) on *CCND1* knock-out brain sections, at E14.5. M-R) Images of the thalamus of wild-type embryonic brains at E14.5 stained by immunofluorescence with antibodies against Ccnd1 (M,P), βIII tubulin (βIIITub, N), or nestin (Q) and their corresponding merges (O, Ccnd1 in green;βIII tubulin in red; R, Ccnd1 in green;nestin in red). The progenitor cell niche is at the bottom of the images. Arrowheads in R point to the co-staining Ccnd1/nestin at the distal part of RGPs (in yellow). S-V) Images of the distal end of RGP in the basal ganglia of wild-type embryonic brains at E14.5, stained by immunofluorescence with antibodies against β1-integrin (S) and Ccnd1 (T) and the merge (U, Ccnd1 in green;β1-integrin in red). Dashed square in U indicate the position of panel V. V) Magnified image of the region shown in U. Arrows indicate partial overlap of Ccnd1 and β1-integrin staining (in yellow) at the tip of the RGP. Scale bars: 300 μm (A), 100 μm (J), 50 μm (M,S). Abbreviations: bg, basal ganglia; ctx, cortex; th, thalamus; v, ventricle.

### *CCND1* knock-out embryos display normal proliferation but abnormal distribution of Tbr2+ and Ctip2+ cells in the developing cortex

We addressed the in vivo relevance of Ccnd1 during brain development by studying *CCND1* knock-out embryos. Given the role of Ccnd1 in the control of cell cycle, we first assessed proliferation defects in the developing cortex using EdU incorporation or PH3 antibodies. Surprisingly, the distribution and the density of EdU+ or PH3+ cells in the cortex from *CCND1* knock-out embryos was not affected, indicating that Ccnd1 is dispensable for proliferation of cortical progenitors (Fig. 2A-F). Next, we studied the structure of the embryonic and postnatal cortex of these animals with several markers including Tbr2 or Ctip2, which label intermediate progenitor cells in the SVZ during development and deep-layer neurons of the mature cortex, respectively. At embryonic stages 16.5 and 18.5 (E16.5 and E18.5), we found that *CCND1* knock-outs exhibited Tbr2+ cells scattered in upper cortical positions, outside of the SVZ (Fig. 2G-L). Likewise, at postnatal stage 8 (P8), we observed that some Ctip2+ neurons, which in control animals are restricted to cortical layer V, were ectopically located in upper and deeper positions (Fig. 2M-O). Other markers, such as Tbr1 and Cux1 were unaffected in these knock-out animals (data not shown). Altogether, these results indicate that Ccnd1 is required for proper cortical layering independently of its role on cell proliferation and therefore suggest a different mechanism of action of Ccnd1 during brain development.

**Figure 2.**
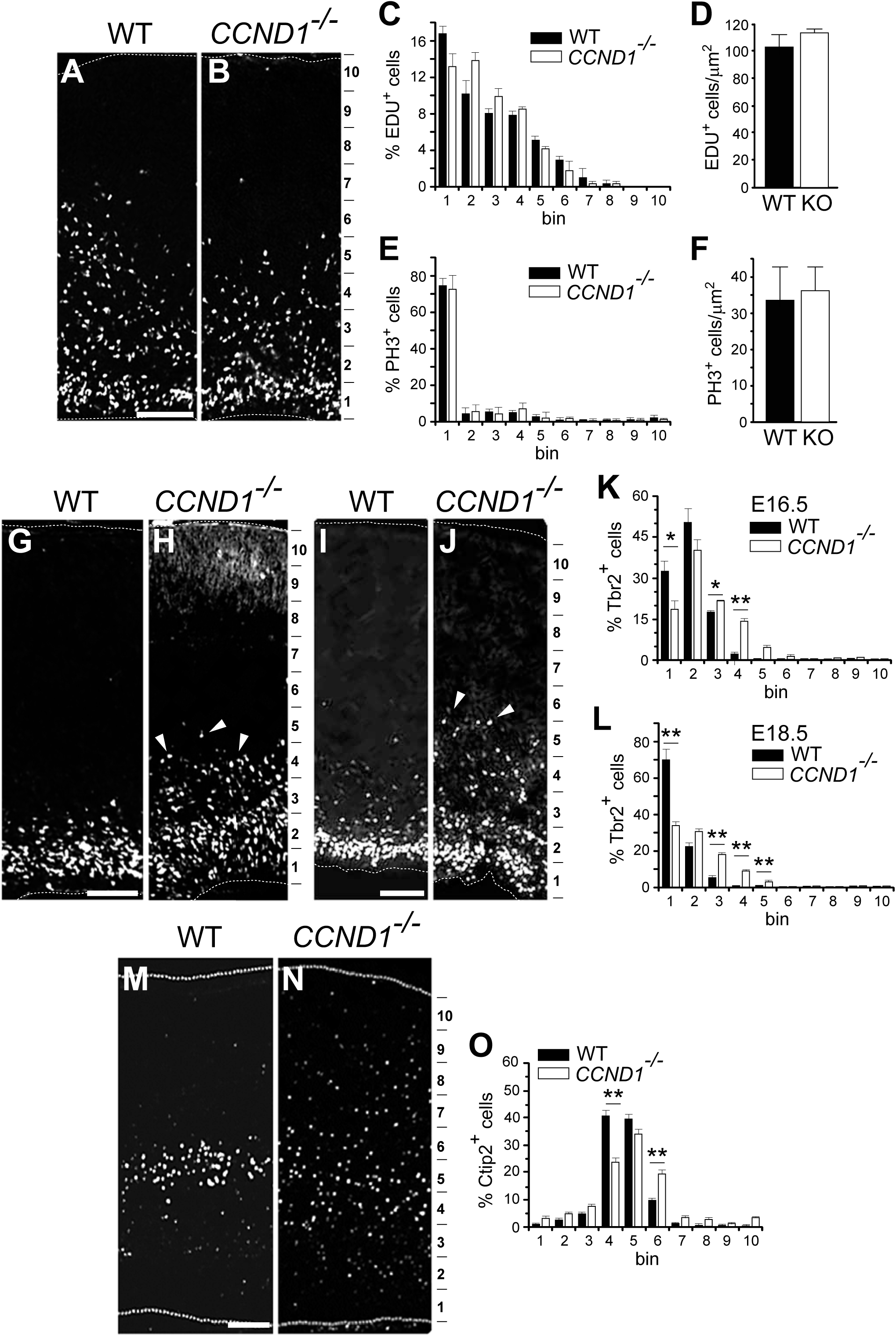
*CCND1* knock-out animals display cortical layering abnormalities but not proliferation defects. A,B) Representative images of cortical sections from wild-type (WT) or *CCND1* knock-out embryos (E14.5), obtained from pregnant females injected for 1 h with EDU. The 10 bin division of the cortical wall used for quantification is indicated on the right. C) Quantification of the percentage of EDU+ cells in each bin along the cortical wall of the sections shown in A,B (black bars, WT; white bars, *CCND1* knock-out). Values are expressed as mean ± SEM (n=3). No statistical differences were observed (two-way ANOVA). D) Comparison of the density of EDU+ cells in cortical sections (E14.5) from wild-type (WT, black bar) or *CCND1* knock-out embryos (KO, white bar). Values are expressed as mean ± SEM (n=3). No statistical differences were observed (*t*-Student; n=3 animals per condition). E) Quantification of the percentage of PH3+ cells in each bin along the cortical wall of the sections stained with anti-PH3 antibodies (black bars, WT; white bars, *CCND1* knock-out). Values are expressed as mean ± SEM (n=3). No statistical differences were observed (two-way ANOVA; n=3 animals per condition). F) Comparison of the density of PH3+ cells in cortical sections (E14.5) from wild-type (WT, black bar) or *CCND1* knock-out embryos (KO, white bar). Values are expressed as mean ± SEM (n=3). No statistical differences were observed (*t*-Student; n=3 animals per condition). G-J) Representative images of cortical sections from wild-type (WT) or *CCND1* knock-out embryos at E16.5 (G,H) or E18.5 (I,J) stained with Tbr2 antibodies. The 10 bin division of the cortical wall used for quantification is indicated on the right. Arrowheads in H and J point to Tbr2+ cells outside the SVZ. K,L) Quantification of the percentage of Tbr2+ cells in each bin along the cortical wall of sections stained in G-J, as indicated (black bars, WT; white bars, *CCND1* knock-out). Values are expressed as mean ± SEM (n=4). Significance was determined by two-way ANOVA and Sidak’s multiple comparison (*p<0.05;**p<0.01). M,N) Representative images of cortical sections from wild-type (WT) or *CCND1* knock-out postnatal brains (P8), stained with anti-Ctip2 antibodies. The 10 bin division of the cortical wall used for quantification is indicated on the right. O) Quantification of the percentage of Ctip2+ cells in each bin along the cortical wall of sections stained in M,N (black bars, WT; white bars, *CCND1* knock-out). Values are expressed as mean ± SEM (n=3). Significance was determined by two-way ANOVA and Sidak’s multiple comparison (**p<0.01). Scale bars: 100 μm.

### Neurons expressing a cytoplasmic dominant negative Ccnd1 (Ccnd1^CAAXK112E^) reproduce some of the layering defects observed in the *CCND1* knock-out embryos

We and others have reported additional functions of Ccnd1 besides cell cycle control, including the regulation of cytoplasmic targets (Zhong et al., 2010; Fusté et al., 2016a; Hydbring et al., 2016). Since Ccnd1 in the RGC process was observed adjacent to the plasma membrane, overlapping with β1-integrin, we investigated if in this localization Ccnd1 had a specific role during cortex development. For that we constructed a version of the protein with a preferred localization in cytoplasmic membranes. This was achieved by adding the CAAX box of KRas, including a polybasic domain adjacent to the CAAX motif, to the C-terminus of Ccnd1 (Ccnd1^CAAX^) that allows its prenylation and tethering mainly to the plasma membrane (Gao et al., 2009; Hancock et al., 2003). We have previously validated this mutant and observed that Ccnd1^CAAX^ was mostly localized at the plasma membrane and cytoplasm and was barely present in the nucleus (Fusté et al., 2016b). Similarly, in dissociated cortical neurons, transfected Ccnd1^CAAX^ shows a cytoplasmic expression, fulfilling the neuron processes and excluding the nucleus (see Extended Data Fig. 5-1,L). Ccnd1^CAAX^ was then subcloned into the pCAGIG vector that contains an IRES EGFP reporter gene for tracking of the electroporated cells and for visualization of their morphology. The construct was then electroporated in embryonic mouse brains by in utero electroporation (IUE) and compared to an empty vector control (pCAGIGempty). We also included an additional control, a cytoplasmic dominant negative Ccnd1 (Ccnd1^CAAXK112E^), in which the lysine 112 (K112) was substituted by a glutamic acid (E) rendering a protein that while still binding Cdks is unable to trigger their kinase activity (Landis et al., 2006). We IUE these constructs at E13.5 and studied the structure and laminar organization of the embryonic and postnatal cortex of the electroporated animals. First, we studied the distribution of the Tbr2+;EGFP+ double positive cells in the developing cortex at E16.5, three days after electroporation. In the empty vector control the percentage of Tbr2+;EGFP+ outside the SVZ among the total EGFP+ cells was very low, as expected (Fig. 3A-D,M). In the condition where Ccnd1^CAAX^ was electroporated, no significant differences were observed (Fig. 3E-H,M). In contrast, electroporation of the dominant negative Ccnd1^CAAXK112E^ triggered a massive increase of the percentage of Tbr2+;EGFP+ cells outside the SVZ, including the intermediate zone (IZ) and upper regions of the CP (Fig. 3I-L,M). These ectopic Tbr2+ cells were negative for proliferation markers such as PH3, indicating that they are not proliferating cells (data not shown). Interestingly, this is reminiscent of the abnormal distribution of Tbr2+ cells that we observed in the IZ of *CCND1* knock-out embryos (Fig. 2G-L). However, the ectopic Tbr2+ cells found within the CP were not observed in the *CCND1* knock-out mutants, suggesting that perhaps its dominant negative effect sequesters more Cdk effectors than those affected by *CCND1* deletion, causing a stronger effect. These results suggest that the cytoplasmic fraction of Ccnd1 is required for the proper layering of the embryonic cortex. In agreement with this interpretation, we also studied the distribution of Ctip2+;EGFP+ double positive cells at P8 in Ccnd1^CAAXK112E^ electroporated brains, and found that many mispositioned outside of layer V (data not shown) resembling what we observed in the *CCND1* knock-out postnatal brains (Fig. 2M-O). In conclusion, our results indicate that cytoplasmic Ccnd1 plays an important role during cortex development and layer organization.

**Figure 3.**
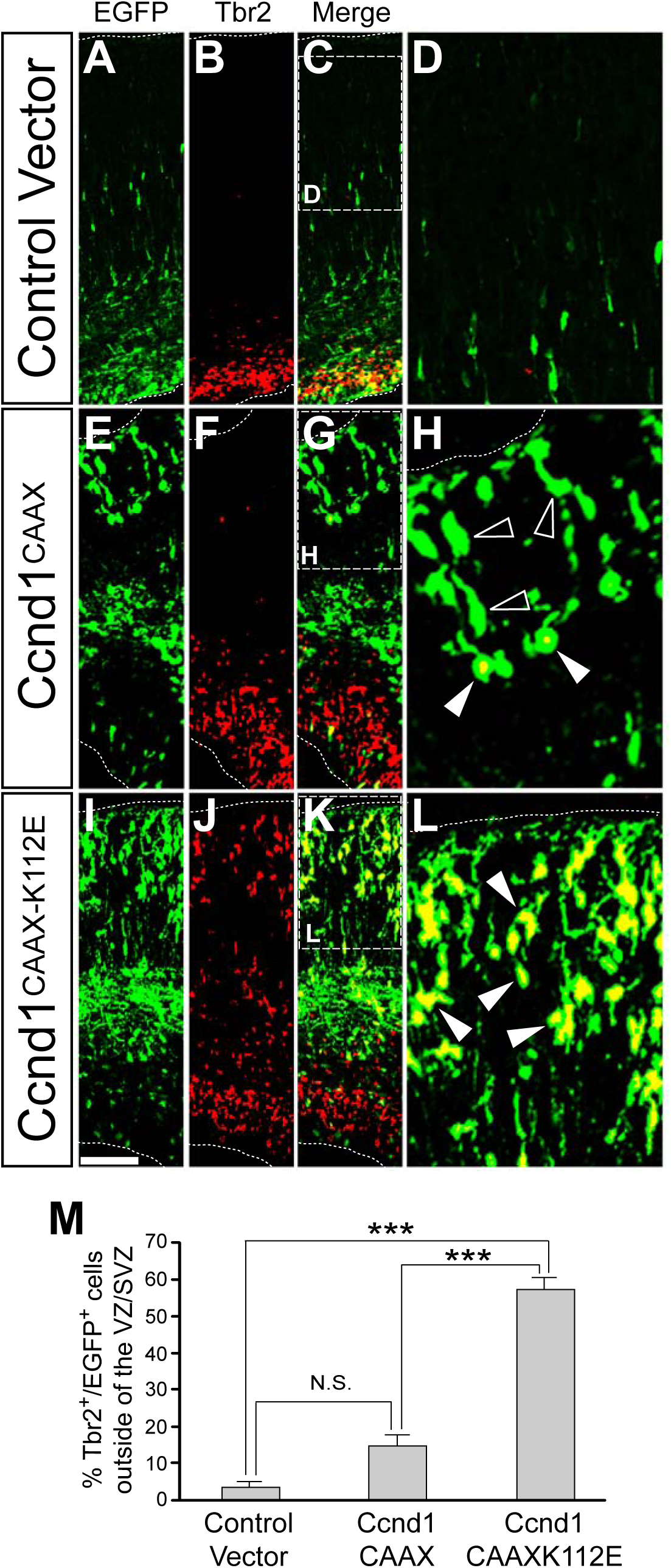
Expression of Ccnd1^CAAXK112E^ affects the distribution of Tbr2+ cells in the developing cortex. Wild-type embryos were IUE at E13.5 with the indicated constructs in pCAGIG vector, and fixed at E16.5: empty vector (A-D), Ccnd1^CAAX^ (E-H) or Ccnd1^CAAXK112E^ (I-L) A-C, E-G, I-K) Overview images of the electroporated cortex with the indicated constructs stained by immunofluorescence with antibodies against EGFP (A,E,I) or Tbr2 (B,F,J). Merged images are shown in C,G,K (EGFP in green; Tbr2 in red). Dashed squares in C, G and K indicate the position of panels D, H and L, respectively. D,H,L) Magnified images of the CP indicated in C,G,K of the merged EGFP/Tbr2 double staining. Empty arrowheads point to EGFP-only stained cells; solid arrowheads point to cells displaying both markers. M) Quantification of the percentage of double EGFP/Tbr2+ cells outside of the VZ/SVZ among the total EGFP+ cells in the cortex of the electroporated embryos with the indicated plasmids. Values are expressed as mean ± SEM (n=5). Significance was determined by one-way ANOVA and Tukey-HSD post-hoc test (***p<0.01; N.S., non-statistically significant difference). Scale bar: 100 μm.

### Cytoplasmic, membrane-associated, Ccnd1 affects the morphology of progenitor cells and of migrating neurons and affects cortical neuron distribution

To understand the base of these layering defects, we performed shorter IUE time experiments in which embryos were collected 12 h after electroporation at E13.5. As expected, after 12 h most of the electroporated cells with pCAGIGempty were progenitors in the VZ/SVZ, with many RGPs extending towards the basal surface (Fig. 4A). In contrast, very few RGPs were observed when Ccnd1^CAAX^ was electroporated (Fig. 4B). On the other hand, brains electroporated with the dominant negative Ccnd1^CAAXK112E^ displayed many more and longer RGPs than empty vector control (Fig. 4C). Since the electroporation time was very short, we assumed that Ccnd1^CAAX^ accelerated the retraction of the preexisting RGP while Ccnd1^CAAXK112E^ slower this process, indicating that proper detachment of the RGP from the meningeal basement membrane during cortex development requires membrane-associated cytoplasmic Ccnd1/Cdk activity. Three days after electroporation (E16.5) we observed that the distribution of both Ccnd1^CAAX^ and Ccnd1^CAAXK112E^ electroporated cells shifted significantly from deeper to upper regions of the cortex, compared to electroporated controls (Fig. 4D). The cells found in the highCP adopted different morphologies and they were multipolar, unipolar/bipolar or were attached to the basement membrane (ABM). In the empty vector control condition, most cells in the highCP displayed unipolar/bipolar morphology or were ABM (Fig. 4E-G). However, Ccnd1^CAAX^ expressing cells displayed a robust increase of multipolar cells in the CP compared to cells electroporated with the empty vector control or with the Ccnd1^CAAXK112E^ (Fig. 4E-G). By contrast, electroporation with Ccnd1^CAAXK112E^ showed a significant increase of ABM cells in the CP (Fig. 4E-G). In summary, these results demonstrate that changes of cytoplasmic, membrane-bound, Ccnd1 expression modulate the morphology of the RGP of progenitor cells and migrating neurons in vivo, during cortex development.

**Figure 4.**
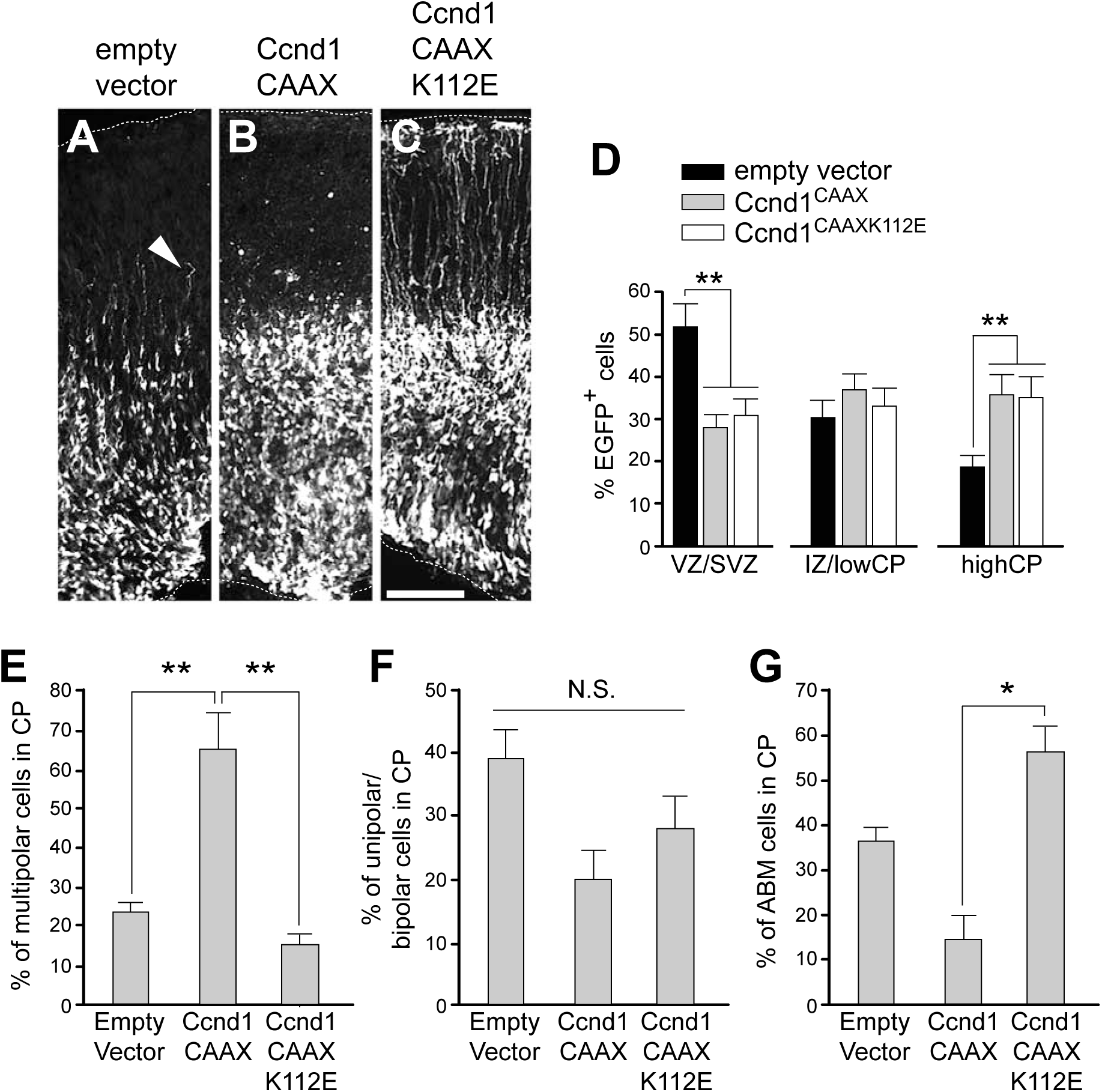
Cytoplasmic Ccnd1 affects RGP and neuron morphology in vivo. A-C) Representative images of cortical sections from wild-type mouse embryos (E13.5) IUE with the indicated constructs in pCAGIG vector. After 12 hrs, the brains were fixed and analyzed by immunofluorescence with an anti-EGFP specific antibody. Arrowhead in A points to the RGPs. Scale bar: 100 μm. D) Quantification of the percentage of EGFP+ cells along the cortical wall divided in three regions VZ/SVZ, IZ/lowCP and highCP of embryos IUE with empty vector (black bars), Ccnd1^CAAX^ (grey bars) or Ccnd1^CAAXK112E^ (white bars) at E13.5 and fixed at E16.5. Values are expressed as mean ± SEM (n=4). Significance in each region was determined by one-way ANOVA and Tukey-HSD post-hoc test (**p<0.01). E-G) Quantification of the different cell morphologies of the EGFP+ cells found in the highCP: multipolar (E), unipolar (F) or attached to the basement membrane (ABM, G) of embryos IUE with the indicated vectors at E13.5 and fixed at E16.5. Values are expressed as mean ± SEM (n=4). Significance in each region was determined by one-way ANOVA and Tukey-HSD post-hoc test (**p<0.01); N.S., non-statistically significant difference.

### Cytoplasmic Ccnd1 promotes neurite and axon outgrowth and increases neurite number in primary cortical neurons

In order to assess more in detail the morphological effects of cytoplasmic Ccnd1 observed in vivo by IUE, we used dissociated cortical neurons from embryonic mouse brains. In these cultures, *CCND1* expression declines quickly, being almost undetectable after 3DIV (Extended Data Fig. 5-1,A). In fact, experimental re-expression of wild-type Ccnd1 induced cell death (Extended Data Fig. 5-1,B-F), an effect that has been widely reported in neurons (Sumrejkanchanakij et al., 2003). We then co-transfected these neurons with Ccnd1^CAAX^ or Ccnd1^CAAXK112E^ together with a plasmid encoding EGFP, as a reporter. Interestingly, overexpression of any of the two cytoplasmic, membrane-bound, Ccnd1 constructs did not trigger cell death and the cultures were viable after 4DIV (Extended Data Fig. 5-1,G-K and data not shown). Instead, Ccnd1^CAAX^ significantly increased the length of the main axon, the total length of neurites and the number of neurites per cell compared to the empty vector control (Fig. 5A,B,D,F-H). These effects were not observed in cultures transfected with Ccnd1^CAAXK112E^ indicating that Cdk activity is necessary (Fig. 5C,E,F-H). Consistent with these observations, the transfection of a similar mutant of Cdk4 (Cdk4^CAAX^) was sufficient to trigger similar morphological changes (Extended Data Fig. 5-1,M,N). In addition, we performed loss-of-function experiments to address the role of Ccnd1 in neuron morphology at 1DIV. First, we transfected neurons with a specific *CCND1* shRNA (Extended Data Fig. 5-1,O) and observed the opposite effect to Ccnd1^CAAX^ overexpression, i.e. significant reductions of axon and neurite lengths, and of the number of neurites per cell compared to a scramble control (Fig. 5I-K). Second, we used cultures from *CCND1* knock-out embryos and observed similar results as in the knock-down approach, a significant reduction of axon and neurite length compared to control littermates (Extended Data Fig. 5-1,P-R). Altogether, these results reveal that Ccnd1 modulates neuron morphology in vitro and that the cytoplasmic, membrane-bound, fraction of the protein, through the activation of Cdks, is the major effector of this function.

**Figure 5.**
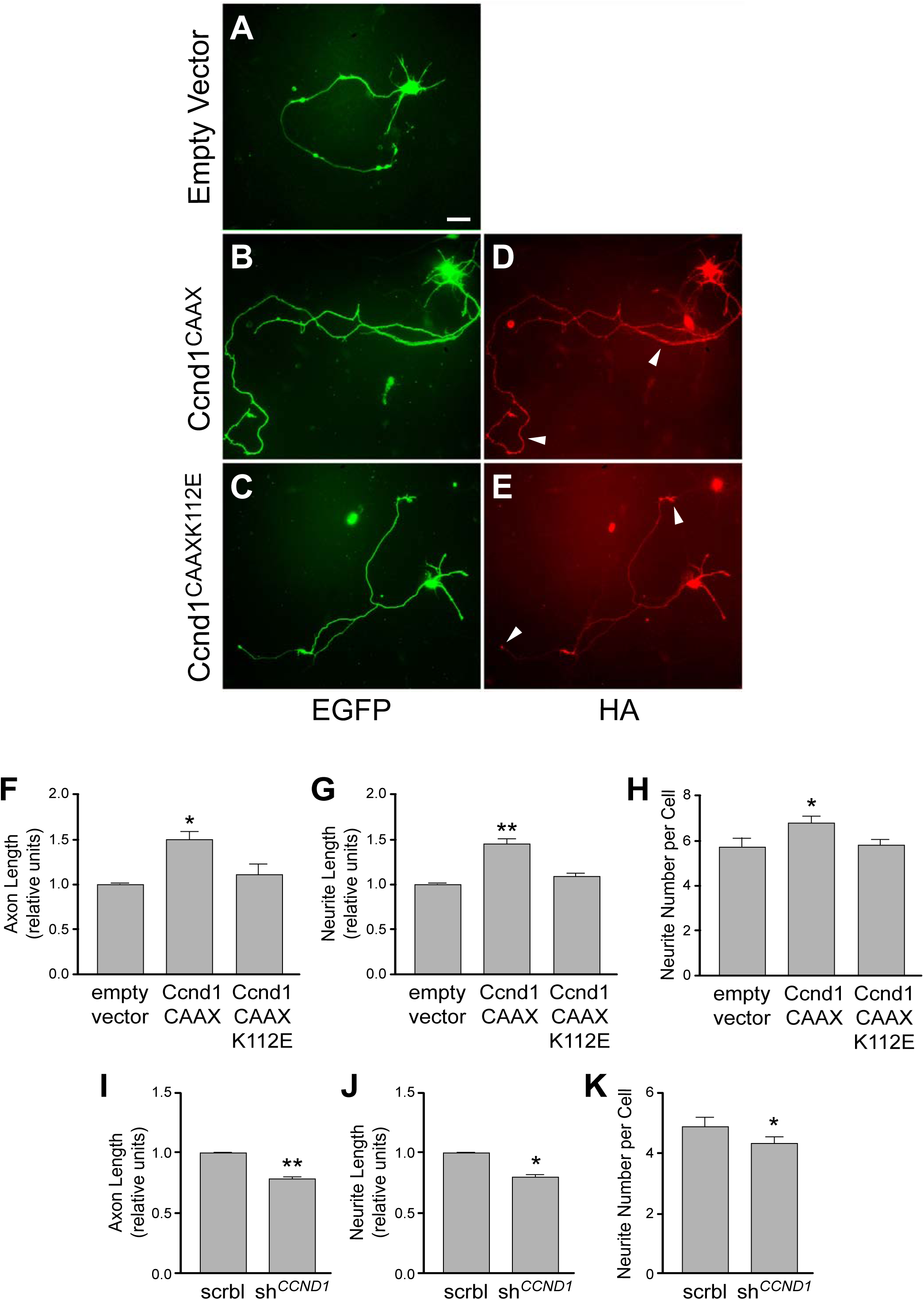
Cytoplasmic Ccnd1 affects neuron morphology in vitro in a Cdk-dependent manner. A-E) Representative images of mouse cortical neuron cultures co-electroporated before seeding with the indicated constructs with an HA tag in pcDNA3 (empty vector as a negative control), and a plasmid encoding EGFP. After 4DIV, cultures were fixed and analyzed by immunofluorescence with antibodies against EGFP (green) and HA (red). Arrowheads in D,E indicate the cytoplasmic expression of Ccnd1^CAAX^ and Ccnd1^CAAXK112E^, respectively. Scale bar, 10 μm. F-H) Neurons from experiments A-E were analyzed for the relative axon length (F), the relative neurite length (G) and the neurite number per neuron (H) using the NeuronJ plugin of ImageJ. Values are expressed as mean ± SEM (n=3). Significance was determined by one-way ANOVA and Tukey-HSD post-hoc test (*p<0.05;**p<0.01). I-K) Cultures of mouse cortical neurons were co-transfected before seeding with plasmids encoding either scramble (scrbl) or *CCND1* shRNA (sh*^CCND1^*) and a plasmid encoding EGFP, as indicated. After 1 DIV, cultures were fixed and analyzed by immunofluorescence with antibodies against EGFP. Relative axon length (I), relative neurite length (J) and the neurite number per neuron (K) were analyzed using the NeuronJ plugin of ImageJ. Values are expressed as mean ± SEM (n=3). Significance was determined by one-way ANOVA and Tukey-HSD post-hoc test (*p<0.05;**p<0.01).

### Cytoplasmic effects of Ccnd1 are mediated by the integrin signaling complex effector paxillin

We have previously shown that Ccnd1 associates with paxillin at the plasma membrane and regulates fibroblast and tumor cell migration and cell-matrix adhesion through Cdk4 and the phosphorylation of paxillin, a member of the signaling complex of integrins in focal adhesions (Fusté et al., 2016a). We characterized the serine residues 83 and 178 as the major phosphorylation sites of Ccnd1/Cdk complexes (Fusté et al., 2016a). We therefore addressed the possibility that paxillin mediates the cytoplasmic effects of Ccnd1 in neurons at the plasma membrane. First, we observed that paxillin phosphorylation at S83 and S178 rapidly decreased in culture (Fig. 6A-C). Interestingly, this response correlated with the reduction of Ccnd1 levels (Extended Data Fig. 5-1,A), suggesting that paxillin might be a target of Ccnd1/Cdk complexes also in neurons. This was confirmed by using palbociclib, a Cdk4/6 specific inhibitor, which completely abolished paxillin phosphorylation at S83 in slices of mouse hippocampi (Fig. 6D). To interfere with the Ccnd1/Cdk-dependent paxillin phosphorylation within the cells we used a phosphorylation mutant where the two serine residues 83 and 178 targeted by Ccnd1/Cdks were substituted by alanine residues (paxillin^S83A;S178A^; Fusté et al., 2016a). When this mutant was transfected in cortical neurons, both neurite and axon length were significantly reduced (Fig. 6E,F). This effect was similar to that observed using *CCND1* knock-out or knock-down cultures (Fig. 5I-K and Extended Data Fig. 5-1,Q,R) and the opposite of cytoplasmic Ccnd1 overexpression (Fig. 5F-H). Altogether, these results are consistent with paxillin being an important effector of membrane-associated Ccnd1/Cdk complexes for controlling neuron morphology, in vitro. We finally addressed if paxillin phosphorylation by Ccnd1/Cdks could also be relevant in vivo during nervous system development. For this, we IUE paxillin^S83A;S178A^ in cortical progenitors at E13.5 and compared the morphology of the electroporated cells with empty vector or wild-type paxillin, 12 h later. This short time was selected to visualize exclusively the preexisting RGPs. We observed that wild-type paxillin caused a significant detachment and retraction of the RGPs from the meningeal basement membrane as the number of RGPs attached to the basement membrane and their length was dramatically reduced compared to empty vector control (Fig. 6G-I). This effect was not observed when paxillin^S83A;S178A^ was electroporated (Fig. 6G-I). Instead, compared to empty vector control, paxillin^S83A;S178A^ displayed an increase of the number of RGPs bound to the meningeal basement membrane (although not statistically significant) and a significant increase of RGP length (Fig. 6H,I). Taking in consideration these observations and the results from Fig. 4A-C, we conclude that both Ccnd1 and paxillin are required for the correct detachment of the RGP from the basement membrane, in vivo, and that both are probably part of the same signaling pathway in this context in which Ccnd1/Cdk complex phosphorylates paxillin at the plasma membrane.

**Figure 6.**
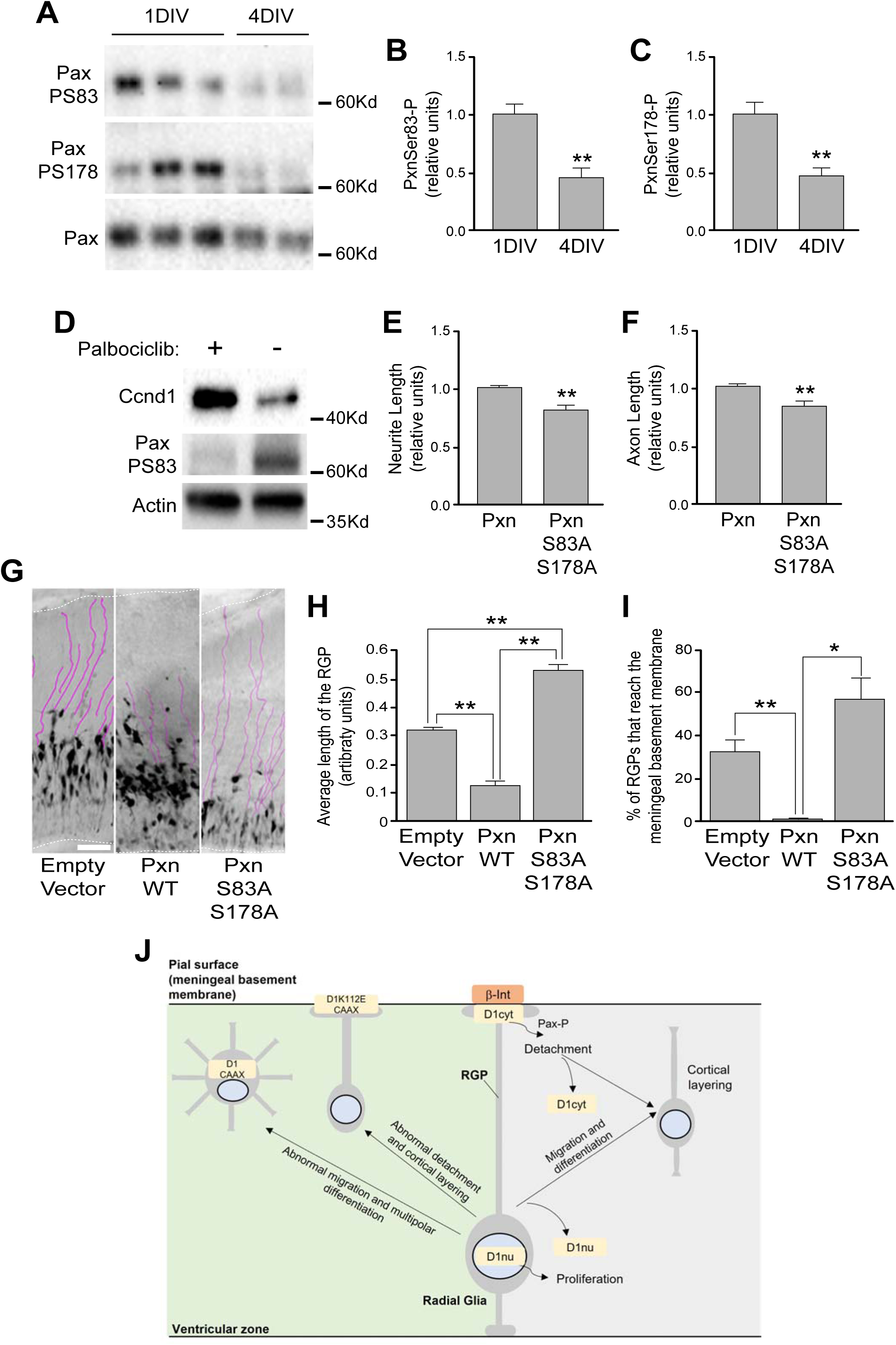
Phosphorylation of paxillin is mediated by Ccnd1/Cdk and is required for neuron differentiation, in vitro and for RGP detachment from the meningeal basement membrane in vivo. A-C) Western-blot analysis (A) and quantification (B,C) of the phosphorylation of paxillin on protein extracts from cortical neuron cultures after 1 or 4 DIV with specific phospho antibodies (phospho-serine83, upper panel; phosphor-serine178, middle panel). Lower panel in A show the total levels of paxillin. Quantification of the phosphorylation of paxillin was corrected by the total paxillin levels and values are expressed as mean ± SEM (n=3). Significance was determined by *t*-Student (**p<0.01). D) Representative Western-blot analysis of the phosphorylation of paxillin (phospho-serine 83, middle panel) on hippocampal slices in the presence of the Cdk4/6 inhibitor palbociclib (2.5 μM), as indicated. Upper and lower panels show the levels of Ccnd1 and actin, respectively. E-F) Cortical neuron cultures were co-electroporated as in Figure 6 (A-H) with wild-type paxillin and the non-phosphorylatable paxillin (Pxn and PxnS83AS178A, respectively) and a plasmid encoding EGFP and were analyzed after 1 DIV for the relative neurite (E) and axon (F) length. Values are expressed as mean ± SEM (n=3). Significance was determined by *t*-Student (**p<0.01). G) Representative images of cortexes IUE at E13.5 with paxillin wild-type (Pxn WT), the non-phosphorylatable paxillin (PxnS83AS178A) or the empty vector control (pCAGIG), as indicated and analyzed at E14.5. RGP were highlighted by purple lines. Scale bar, 100 μm. H,I) Quantification of the average RGP length (H) or the percentage of RGP attached to the meningeal basement membrane in the conditions described in panel G. Values are expressed as mean ± SEM (n=3). Significance was determined by one-way ANOVA and Tukey-HSD post-hoc test (*p<0.05;**p<0.01). J) Summary of our results. Right part (light grey): during normal cortex development Ccnd1 needs to be downregulated both from the nucleus (D1nu) and from the RGP (D1cyt) of cortical progenitors to stop cell division and allow migration and differentiation and to detach from the meningeal basement membrane, respectively. In the later case, phosphorylation of paxillin (Pax-P) by cytoplasmic Ccnd1/Cdk complexes is required. Left part (light green): expression of cytoplasmic Ccnd1 (D1CAAX) triggers abnormal migration and multipolar shape of differentiating neurons while expression of a dominant negative Ccnd1, with cytoplasmic localization (D1K112ECAAX), triggers abnormal detachment of RGP from the meningeal basement membrane and layering defects.

## Discussion

During nervous system development, the molecular mechanisms controlling proliferation are tightly regulated to achieve the correct timing for neuron differentiation. In the present work we have shown that Ccnd1 plays key functions during cortex development independent of its well-established role on cell cycle regulation. Instead, we have shown that Ccnd1 is also localized in the cytoplasm and associated to the plasma membrane of the RGPs of cortical neuron progenitors. Our results suggest that in this localization Ccnd1 regulates meningeal basement membrane adhesion through paxillin, a downstream effector of integrins in focal adhesions. In addition, we show that alteration of cytoplasmic Ccnd1 levels affect cortical layering and neuron morphology in vivo and in vitro. A summary of our results is shown in Fig. 6J.

As it was reported previously, we found that the brain of *CCND1* knock-out animals developed normally, without gross morphological defects and no alteration of specific proliferation markers (Glickstein et al., 2009). In fact, just mild proliferation defects were described at P0 in the granule progenitor cells leading to a reduction of postnatal cerebellar size (Pogoriler et al., 2006). Ccnd2 seems to have a more prominent role in the proliferation of cortical precursors in vivo as its genetic ablation causes G1 lengthening, premature cell cycle exit and faster neuronal differentiation, leading to microcephaly and thinner cortical wall (Glickstein et al., 2009). The *CCND1* loss-of-function data contrasts with the effects of Ccnd1 overexpression in cortical progenitors. Electroporation of Ccnd1/Cdk4, Ccnd1 or Ccne1 shortens G1, promoting the expansion of progenitor cells and delaying neurogenesis (Lange et al., 2009; Pilaz et al., 2009). Similar results were observed in neural stem cells of the adult hippocampus and olfactory bulb (Artegiani et al., 2011; Bragado Alonso et al., 2019). Previous studies have suggested redundant functionality among Ccnds, in vivo (Kenney and Rowitch, 2000; Ciemerych et al., 2002; Satyanarayana and Kaldis, 2009). Both Ccnd2 and Ccnd1 display partial overlapping expression in the developing cortex and Ccnd2 can indeed replace the function of Ccnd1 in cortical proliferation (Glickstein et al., 2007; Glickstein et al., 2009).

Besides cell cycle control, Ccnd1 exhibits additional functions in multiple tissues and cell types suggesting a more intricate mechanism of action (Hydbring et al., 2016). For instance, in granule progenitor cells, Ccnd1/Cdk4 complex phosphorylates the transcription factor ATOH1, preventing its degradation, which is essential for maintaining their immature state (Miyashita et al., 2021; Flora et al., 2009). By contrast, in cortical neuron progenitors Ccnd1 promotes lineage-commitment and differentiation while Ccnb1/2 favour self-renewal of radial glial cells (Hagey et al., 2020). In line with this last study, we show a graded expression of Ccnd1 in the developing cortex, from ventro/lateral-high to dorso/medial-low, that has been reported to be controlled by the transcription factor Sp8 and related to neuronal differentiation (Borello et al., 2018). Pro-neurogenic functions of Ccnd1 were previously reported. For example, Ccnd1 expression, but not Ccnd2, is maintained during the initial phase of motoneuron differentiation and stimulates neurogenesis by a cell cycle-independent mechanism involving the pro-neurogenic transcription factor Hes6 (Lukaszewicz and Anderson, 2011). In addition, in PC12 cells the neurotrophic factor NGF activates Ccnd1 expression, which is necessary for neurite outgrowth (Marampon et al., 2008). Consistently, we show that ablation or downregulation of *CCND1* in mouse cortical neurons cause a significant reduction of neurite complexity. This could be explained by the regulation of pro-neurogenic transcription factors by Ccnd1 (Lukaszewicz and Anderson, 2011). However, our results point to an alternative, not mutually exclusive, interpretation. We suggest that cytoplasmic, membrane associated, Ccnd1 is required for neuron differentiation as the overexpression of Ccnd1^CAAX^ enhances neuron complexity while Ccnd1^CAAXK112E^ has no effect, indicating the requirement of Cdk activity in this effect. Thus, we provide a novel cell cycle-independent function of Ccnd1/Cdk complex that requires cytoplasmic localization.

Cytoplasmic localization of Ccnd1 in postmitotic hippocampal and cortical neurons and neuroblastoma cells was previously observed. This was first proposed as a mechanism to prevent apoptosis and/or for cell cycle withdrawal (Sumrejkanchanakij et al., 2003; Sumrejkanchanakij et al., 2006; Schmetsdorf et al., 2005). However, we and others have provided evidence in cancer cells and embryonic fibroblasts that cytoplasmic Ccnd1 can be found as well associated to the plasma membrane and has an active role in controlling cell adhesion or motility with different intracellular mechanisms involved (see below). In the context of neuronal function, it was recently reported that extracellular vesicles from NGF-differentiated PC12 cells contain Ccnd1, which is necessary to induce neuronal lineage of stem cells (Song et al., 2021). In addition, we have shown that cytoplasmic Ccnd1/Cdk complex controls postmitotic neuronal communication by phosphorylation of the α4 subunit of GABAA receptors (Pedraza et al., 2023). In the present work we show that Ccnd1 is present in the RGP of progenitor cells during cortex development, in vivo, and that is associated to β1-integrin at the plasma membrane of the tip of the basal RGP. This cytoplasmic localization was previously reported for Ccnd2, being important for the regulation of asymmetric cell division during corticogenesis and neuronal differentiation (Glickstein et al., 2007; Tsunekawa et al., 2012). In contrast to our results, this localization was not previously observed for Ccnd1 (Glickstein et al., 2007), probably due to the use of different Ccnd1 antibodies or different tissue preparation protocols. *CCND2* mRNA requires a unique cis-regulatory sequence in its 3’ untranslated region to be transported into the RGP, where is locally translated (Tsunekawa et al., 2012). We have been unable to detect a similar sequence in the *CCND1* mRNA and therefore the mechanisms regulating cytoplasmic localization of Ccnd1 remain to be elucidated.

Our data indicate that cytoplasmic Ccnd1 expression in radial glial cells needs to be tightly regulated to achieve proper cortex formation and layering. Overexpression of cytoplasmic Ccnd1 accelerates their detachment from the meningeal basement membrane and affects the morphology and the positioning of post-mitotic neurons while expression of a dominant negative mutant delays this detachment and cause layering defects of Tbr2+ and Ctip2+ neurons. Some of these layering defects were also observed in *CCND1* knock-out brains, highlighting the importance of cytoplasmic Ccnd1 in vivo. Acute experiments by IUE with the dominant negative mutant show indeed stronger effects than the knock-out, suggesting a broader inhibition of Cdks. Most of the interactors and signalling effectors of cytoplasmic Ccnd1 have been related to cell adhesion at the plasma membrane, including the actin-binding protein filamin A in membrane ruffles (Zhong et al., 2010), the cytoplasmic adapter protein PACSIN 2 in focal adhesions (Meng et al., 2011), the Ral GTPases (Fernandez et al., 2011; Cemeli et al., 2019) and paxillin, a downstream integrin signalling protein in focal adhesions (Fusté et al., 2016a). Here we provide evidence that cytoplasmic Ccnd1 in radial glial cells could be regulating cortex development by integrin signalling through paxillin phosphorylation, adding evidence of the important role of integrin signalling during cortex development (Long and Huttner, 2019). We have shown that Ccnd1 co-localizes with β1-integrin at the plasma membrane of the distal tip of the basal RGP, that Ccnd1 levels correlate with phosphorylation of paxillin in neurons and that this phosphorylation is necessary for neurite extension, in vitro, and for detachment of the RGP from the meningeal membrane, in vivo. Proper detachment of RGP from the meningeal basement membrane has been shown to be essential for cortical layering. For instance, deletion of β1-integrin from progenitor cells, but not from post-mitotic neurons, triggers abnormal development of the glial end feet, disruption of the basal lamina and perturbs cortical layer formation, mainly affecting upper layers (Graus-Porta et al., 2001; Belvindrah et al., 2007). In addition, interference with the adhesion protein TAG-1 (transient axonal glycoprotein-1), also known as contactin-2, provokes basal RGP retraction leading to cortical layering defects including ectopic Tbr2+ progenitors and Ctip2+ cells mislocalization (Okamoto et al., 2013). By contrast, we report an opposite correlation between increased RGP adhesion and layering defects. Although speculative and in need of further investigation, it is possible that exact detachment timing of the RGP from the meningeal basement membrane is essential for cortex layering and that an acceleration or delay will end up in similar layering defects. Maybe longer adhesion shifts the migration type from locomotion to translocation (Nadarajah et al., 2001), explaining the increase in the CP of electroporated neurons bearing the Ccnd1 dominant negative mutant and the concomitant layering defects.

Although the cytoplasmic expression of Ccnd1 was mostly observed in the RGP we cannot rule out that significant amounts of cytoplasmic Ccnd1 could play a role as well in post-mitotic neurons, as previously suggested (Lukaszewicz and Anderson, 2011; Sumrejkanchanakij et al., 2003; Song et al., 2021; Pedraza et al., 2023). Paxillin or β1-integrin knock-out animals display a delay in positioning of upper layer neurons, a cell-autonomous effect in post-mitotic neurons triggered by changes in neuron morphology and by a slower migration (Rashid et al., 2017; Rashid and Olson, 2023). A similar phenotype was observed by ablation of focal adhesion kinase (FAK), which is recruited by paxillin into focal adhesions, reinforcing the role of integrin signalling in post-mitotic neurons during cortex formation (Valiente et al., 2011). According to this data, it is possible that activation of paxillin by Ccnd1 accelerates migration, explaining therefore the accumulation of neurons in the CP upon cytoplasmic Ccnd1 overexpression. Finally, there is also the possibility that other substrates of cytoplasmic Ccnd1/Cdk activity, different from paxillin, could have a role during cortex development.

## Supporting information

Extended Data for Figure 5

## Acknowledgements

We thank the funding agencies Ministerio de Ciencia, Innovación y Universidades, Plan Estatal de Investigación Científica y Técnica y de Innovación (Egea: PGC2018-101910-B-I00 and PID2021-129089NB-I00; Garí/Ferrezuelo: PID2019-104859GB-I00) and the Generalitat de Catalunya, Agència de Gestió d’Ajuts Universitaris i de Recerca (AGAUR), Suport Grups de Recerca SGR2021, 01113 (Garí). We thank members of the Cell Cycle and the Oncogenic and Developmental Signaling groups for fruitful discussions. Special thanks to Sònia Rius for technical support and to Jèssica Pairada and the rest of the animal facility and the tissue culture staff of the University of Lleida, for technical service.

**Extended data supporting Figure 5 (Fig. 5-1).**

A) Relative levels of *CCND1* mRNA in cortical neuron cultures were analyzed by qRT-PCR at the indicated time points. Values are expressed as mean ± SEM (n=3). Significance was determined by one-way ANOVA and Tukey-HSD post-hoc test (***p<0.001).

B-L) Representative images of mouse cortical neuron cultures from E15.5 embryos co-electroporated before seeding with the indicated constructs (HA-tagged). After 4DIV, cultures were fixed and analysed by immunofluorescence with antibodies against HA (red; B,G) and cleaved caspase3 (green; D,I) and counterstained with Hoechst (blue; C,H). The indicated merged images are shown in E,F,J,K,L. Nuclear exclusion and membrane association of Ccnd1^CAAX^ was observed (L). Arrowheads in F and L indicate caspase3+ apoptotic neurons and membrane associated Ccnd1^CAAX^, respectively. Scale bar, 20 μm.

L,M) Mouse cortical neuron cultures co-electroporated before seeding with Cdk4^CAAX^ (HA-tagged) construct in pcDNA3 (empty vector as a negative control), and a plasmid encoding EGFP. After 4DIV, cultures were fixed and analyzed the relative axon length

(L) and the relative neurite length (M) using the NeuronJ plugin of ImageJ. Values are expressed as mean ± SEM (n=3). Significance was determined by *t*-Student (*p<0.05;**p<0.01).

N) Mouse embryonic fibroblasts were infected with lentiviral particles containing control (scrb) or shRNA against Ccnd1 and selected with puromycin at 25 μg/ml for 3 days. Protein extracts were analyzed by Western blot with specific antibodies against Ccnd1 (upper panel) or actin, as loading control (bottom panel). Molecular weight size markers (kDa) are indicated on the right.

O) Protein extracts from the hands of E15.5 embryos from three *CCND1* knock-out (*CCND1^-/-^*) and from three sibling controls (WT) were analyzed by Western-blot with specific antibodies against Ccnd1. Molecular weight size marker (kDa) is indicated on the right.

P,Q) Cultures of mouse cortical neurons from *CCND1* knock-out brains or sibling controls were electroporated before seeding with a plasmid encoding EGFP. After 1 DIV, cultures were fixed and analyzed by immunofluorescence with antibodies against EGFP. Relative neurite length (P) and axon length (Q) were analyzed using the NeuronJ plugin of ImageJ. Values are expressed as mean ± SEM (n=3). Significance was determined by *t*-Student (*p<0.05).

