## Supplementary figures and images for "Cytoplasmic expression of the cell cycle regulator cyclin D1 in radial glial progenitor cells modulates brain cortex development"

### Extended Data for Figure 5

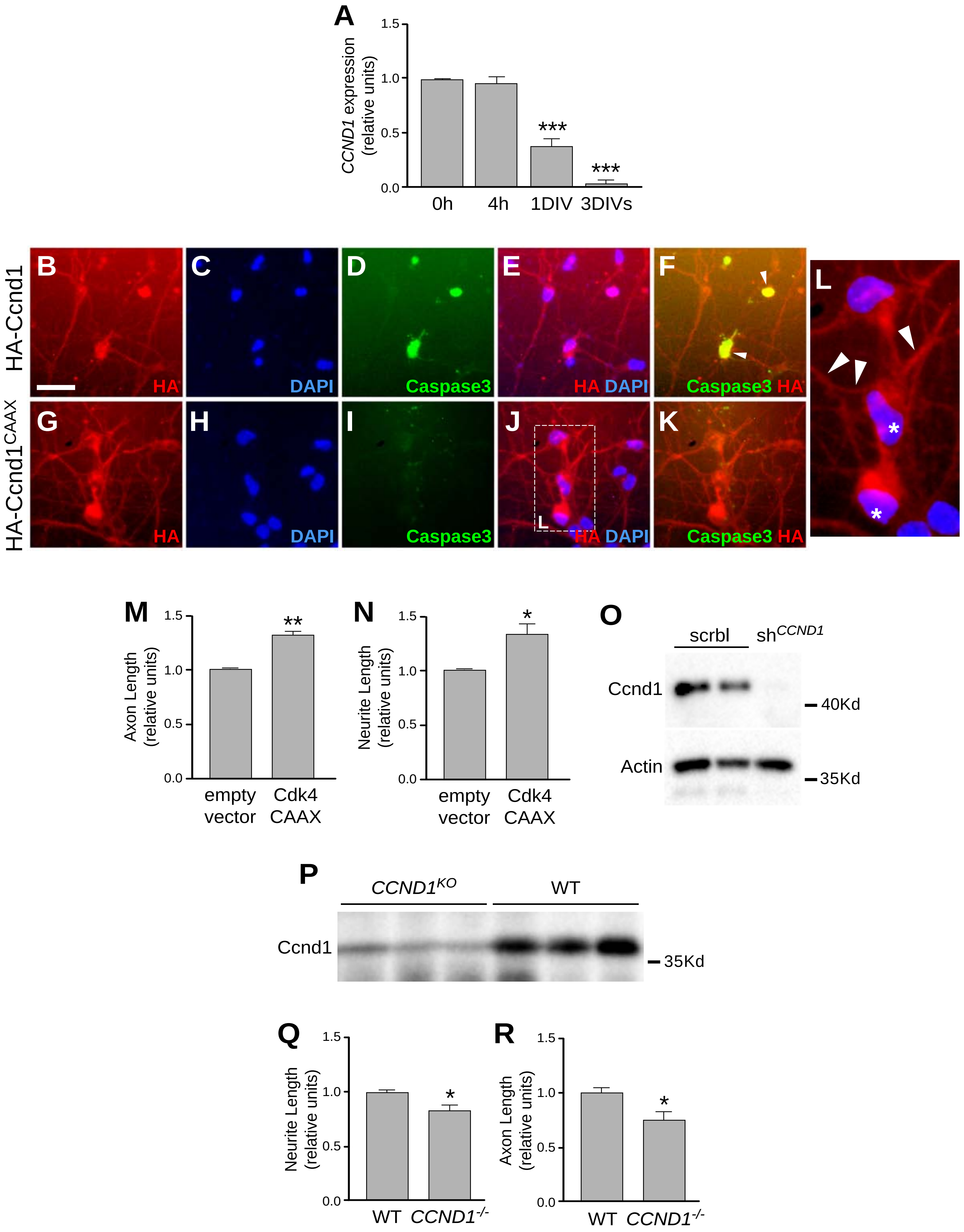


Pedraza et al., Extended Data (Figure 5-1)
